# Fiber enhancement and 3D orientation analysis in label-free two-photon fluorescence microscopy

**DOI:** 10.1101/2022.10.19.512531

**Authors:** Michele Sorelli, Irene Costantini, Leonardo Bocchi, Markus Axer, Francesco Saverio Pavone, Giacomo Mazzamuto

**Author notes:** corresponding author *Email address:* (Irene Costantini).

## Abstract

3D fluorescence microscopy can be profitably used in combination with tissue clearing and myelin labeling techniques for investigating the human brain myeloarchitectonics with sub-micrometric resolution, and for providing detailed histological reference data for the validation of diffusion magnetic resonance imaging (dMRI) tractography. Differently from state-of-the-art polarimetry-based neuroimaging modalities such as 3D polarized light imaging and 3D polarized-sensitive optical coherence tomography, the quantitative histological analysis of cortical and white matter fiber architectures from fluorescence microscopy images requires the estimation of fiber orientations via image processing techniques, as Fourier or structure tensor analysis. However, these can be error-prone as they are not able to detect myelinated fibers in surrounding brain tissue. Here we present a novel image processing pipeline built around a 3D Frangi filter able to enhance fiber structures of varying diameters in tiled brain section reconstructions of arbitrary size, and to produce accurate 3D fiber orientation maps in both grey and white matter regions, while preserving the native resolution of the employed microscopy system. The developed software tool also features the estimation of fiber orientation distribution functions which, in view of their widespread use in the neuroimaging community, may favor a multimodal comparison with brain connectivity data obtained from the aforementioned label-free optical modalities, and the validation of modern dMRI-based tractography. The proposed image processing pipeline was applied to brain samples treated according to a novel label-free preparation technique capable of enhancing the autofluorescence of myelinated fiber axons, thus demonstrating the feasibility of single fiber orientation analyses in fluorescence microscopy without the need for exogenous staining.

## 1. Introduction

The integration of different imaging modalities at micro-, meso- and macroscopic spatial scales is widely recognized as an essential requirement for advancing our current knowledge of the human brain connectional anatomy (Axer et al. (2016); Alimi et al. (2020); Yendiki et al. (2022)). In this regard, the possibility to investigate brain myeloarchitectonics with single fiber resolution over extended tissue volumes represents a compelling research area. The introduction of increased magnetic field strengths and the development of improved theoretical contrast models in diffusion-weighted magnetic resonance imaging (dMRI), fostered by the Human Brain Connectome project (Setsompop et al. (2013)), have recently enabled the quantitative mapping of the human brain connectivity with sub-millimetric resolution, and improved angular accuracy (Tournier (2019); Trampel et al. (2019); Beaujoin et al. (2019); Huang et al. (2021)). The ultra-high b-values available nowadays in clinical and pre-clinical scanners provide increased signal-to-noise ratios and, thus, enhanced spatial resolution; this, in turn, represents a key factor in improving the quality of dMRI-based fiber tractography (Fan et al. (2014); Jones et al. (2020)). However, despite its unique capability to provide an in vivo interrogation of the structural organization and connectivity of the human brain, state-of-the-art dMRI may still inadequately capture the tissue microstructure within voxels containing complex branching or interdigitated fiber architectures (Yendiki et al. (2022)). Therefore, the development of a gold standard method capable of providing reliable multiscale ground-truth datasets for the comprehensive validation of dMRI-based connectivity information is of paramount importance. This has motivated several research groups to apply dMRI to post mortem histological tissue sections in combination with a range of optical modalities, namely 3D polarized light imaging (3D-PLI) (Axer et al. (2011, 2016)), 3D polarized-sensitive optical coherence tomography (3D-PSOCT) (Wang et al. (2018); Jones et al. (2020)), 2D and 3D confocal scanning fluorescence microscopy (Budde and Frank (2012); Khan et al. (2015); Schilling et al. (2018)) and 3D light-sheet fluorescence microscopy (LSFM) (Morawski et al. (2018)), also combining OCT and LSFM (Costantini et al. (2021)), with the aim to identify suitable approaches for validating dMRI-based fiber tractography, and unveiling its potential limitations in brain regions with challenging fiber compositions and geometrical configurations. 3D-PLI and 3D-PSOCT exploit the inherent birefringence of brain tissue, analyzing the change in polarization of light transmitted through unstained samples in order to detect fiber structures and estimate their 3D orientation. On the other hand, fluorescence microscopy generally relies on tissue clearing methods, such as CLARITY (Clear Lipid-exchanged Acrylamide-hybridized Rigid Imaging compatible Tissue hYdrogel) (Chung and Deisseroth (2013); Costantini et al. (2015)), or SWITCH (System-Wide control of Interaction Time and kinetics of CHemicals) (Murray et al. (2015); Costantini et al. (2021); Pesce et al. (2022)) to render samples transparent, and histological stains for generating contrast between fibers and other tissue components. Furthermore, despite offering a superior spatial resolution with respect to polarimetry-based imaging modalities, which enables the targeting of finer microstructural details, in fluorescence microscopy the evaluation of fiber orientations is subject to the application of dedicated image processing techniques, namely Fourier analysis (Budde et al. (2011); Choe et al. (2012)) and structure tensor analysis (Budde and Frank (2012); Khan et al. (2015); Schilling et al. (2018)). However, a relevant limitation of these techniques is the non-specificity of the estimated orientation vectors, as they are not able to distinguish between fibers and surrounding brain tissue. In this regard, Morawski and co-workers (Morawski et al. (2018)) have applied a supervised Random Forest classifier, cascaded with a multi-scale enhancement Frangi filter (Frangi et al. (1998)), for the automatic identification of fiber structures. However, their orientation analysis was limited to the image plane due to the marked anisotropic resolution of the employed light-sheet microscope and, moreover, it relied on the availability of manual segmentations produced by expert neuroanatomists. In this work, we combined the sub-micron resolution and deep imaging capabilities offered by two-photon fluorescence microscopy (TPFM), with the myelin autofluorescence enhancement achieved by a pioneering label-free preparation protocol named MAGIC (Myelin Autofluorescence imaging by Glycerol Induced Contrast enhancement) (Costantini et al. (2021)), for acquiring high-resolution mesoscale reconstructions of the human brain fiber organization. Large volumetric brain tissue images were analyzed by means of a novel image processing pipeline built around a 3D implementation of the Frangi filter, able to further enhance myelinated fiber structures of varying diameters and produce 3D orientation maps preserving the native resolution of the TPFM system. Finally, the analysis pipeline features the generation of orientation distributions functions (ODFs) at arbitrary spatial scales (Bunge (1982)), that may be used in the future to enable a direct comparison with micrometric fiber orientations obtained via 3D-PLI and/or 3D-PSOCT, and to validate the meso- and macro-scale brain connectivity information targeted by state-of-the-art dMRI.

## 2. Materials and Methods

### 2.1. Specimen collection

Tissue sections imaged and analyzed in this study were obtained from a post mortem human brain (male, 87 years). The human brain was acquired in accordance with the ethics committee at the Medical Faculty of the University of Rostock, Germany (#A2016-0083). The body donor provided written informed consent for the general use of post mortem tissue for aims of research and education. The usage is covered by a vote of the ethics committee of the medical faculty of the Heinrich Heine University Düsseldorf (#4863). All methods were carried out in accordance with relevant guidelines and regulations.

### 2.2. MAGIC preparation protocol and two-photon fluorescence microscopy

Human brain samples were preliminarily treated for fluorescence microscopy following the label-free MAGIC preparation technique, developed by Costantini (Costantini et al. (2021)). This method originates from the evidence that glycerol removal from fixed and embedded brain tissue leads to a specific increase in myelin autofluorescence and thus to an enhanced signal-to-background ratio of myelinated axons. In detail, brain tissue was initially fixed with a 4 % paraformaldehyde (PFA) solution for *>*12 weeks. Then, it was first embedded in a 10 % glycerol, 2 % DMSO, 4 % formaldehyde solution and, second, in a 20 % glycerol, 2 % DMSO, 4 % formaldehyde solution for *>*3 weeks. These steps were conducted at a tightly controlled temperature of 4 ^°^C. Afterwards, brain samples were treated with a 2 % dimethyl sulfoxide solution for cryoprotection and immersed in isopentane (−50 ^°^C) for *>*30 minutes. Next, frozen human brains were sliced at a temperature of −30 ^°^C into 60 μm-deep coronal sections using a cryostat microtome (Leica Microsystems, Germany). Finally, brain tissue slices were incubated at ambient temperature in a 0.01 M phosphate buffer saline solution (PBS) for three months. Prior to imaging, sections were coverslipped and mounted in PBS. Brain slices treated with the MAGIC preparation protocol were imaged using a custom TPFM system. As excitation source, the system employs a Chameleon (Coherent, US) tunable mode-locked Ti:Sapphire laser (120 fs pulse width, 90 MHz pulse rate) operating at 800 nm. The laser source is optically coupled with a custom lateral scanning system comprising a pair of galvanometric mirrors (LSKGG4/M, Thorlabs, US), and focused onto the brain tissue slices by means of a tunable 25x objective (LD LCI Plan-Apochromat 25x/0.8 Imm Corr DIC M27, Zeiss, Germany), characterized by a 0.8 numerical aperture and a free working distance of 0.57 mm. The resulting field of view is 450 μm × 450 μm. 482/35 nm and 618/50 nm single-band bandpass emission filters (BrightLine^®^, Semrock, US) were respectively used for detecting myelinated fibers and neuronal bodies. The fluorescence intensity signal was collected by a GaAsP photomultiplier tube (H7422, Hamamatsu Photonics, JP) and digitized with 8-bit precision.

The point spread function (PSF) of the TPFM system was previously characterized in (Costantini et al. (2021)), by imaging 100 nm beads (FluoSpheresTM carboxylate-modified microspheres, yellow-green fluorescent, Thermo Fisher Scientific, US) embedded at a 1:1000 concentration in a PBS gel reproducing the refractive index of the immersion medium employed for the brain slice imaging. The average shape of the PSF was estimated using the Huygens PSF distiller tool (version 19.04, Scientific Volume Imaging, NL), resulting in a measured FWHM of (0.692, 0.692, 2.612) μm along the x, y, and z axes, respectively. No preliminary deconvolution was applied to the TPFM images acquired in this work. The TPFM system features a closed-loop XY stage (U-780 PILine XY Stage System, Physik Instrumente, Germany) enabling the lateral translation of the imaged specimen, and a closed-loop piezoelectric stage (ND72Z2LAQ PI-FOC Objective Scanning System, Physik Instrumente, Germany) for the vertical displacement of the objective. The LabVIEW software controlling the system enables the sequential acquisition of adjacent overlapping 3D tiles, allowing for mesoscale reconstructions of entire brain slices with a maximum lateral resolution of 0.44 μm (corresponding to 1024 px × 1024 px images) and a minimum axial sampling step of 1 μm. A voxel size of 0.88 μm × 0.88 μm × 1 μm and a lateral tile overlap of 40 μm were adopted in the present study.

### 2.3. Frangi filter theory

The Frangi filter produces a specific enhancement of tubular objects in 2D or 3D images, exploiting geometrical information on the local gray level structure derived from the eigenvalue decomposition of Hessian matrices estimated at the pixel level. As detailed in (Frangi et al. (1998)), the structural properties of an image *I* can be characterized within a neighborhood defined by the scale space parameter *σ* through the following second-order Taylor expansion:

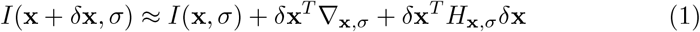

where ∇_**x**,*σ*_ and *H*_**x**,*σ*_ denote the gradient vector and the Hessian matrix of the partial spatial derivates computed in **x** which respectively describe the first- and second-order local structure of the image. In the particular case of a 3D image, the Hessian matrix is defined as:

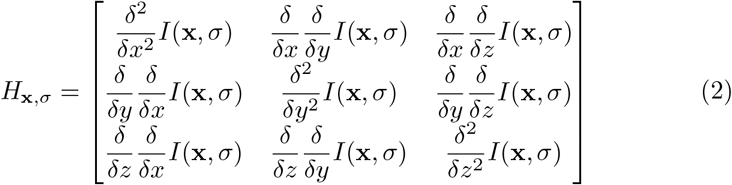

where the scale representation of the image *I* at scale *σ, I*(**x**, *σ*), is generally obtained via a preliminary convolution with a Gaussian kernel of variance *σ*^2^:

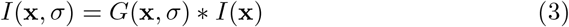

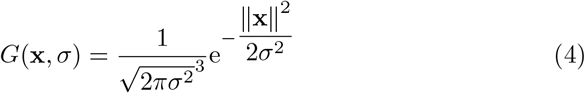

In order to compensate for the progressive attenuation of spatial image derivatives at increasing analysis scales, with the consequential underestimation of *H*_**x**,*σ*_ matrices for larger *σ* values, the Frangi filter makes use of scale-normalized spatial derivatives, that is:

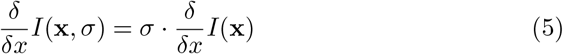

Otherwise, finer image details would be systematically over-enhanced with respect to coarser structures.

The eigenvalue decomposition of the scaled Hessian matrix determines the principal directions of the second-order structure of the image; specifically, the Hessian matrix maps a finite image neighborhood onto a 3D ellipsoid whose semiaxes have a length corresponding to the estimated eigenvalues *λ*_*i*_ and are oriented according to the related eigenvectors **v**_*i*_. This second-order ellipsoid thus provides an intuitive description of the local image behavior, exploited by Frangi et al. to design geometrical features which enable the specific segmentation of tube-like objects.

Indeed, in presence of specific spatial contrast patterns, particular relations must hold between the three eigenvalues (Table 1). Namely, assuming |*λ*_1_| ≤ |*λ*_2_| ≤ |*λ*_3_|, a pixel belonging to a bright 3D tubular structure over a dark background would be ideally characterized by a near-zero *λ*_1_ (the dominant eigenvalue identifying the local direction of minimum intensity variation), and by *λ*_2_ and *λ*_3_ having a large similar magnitude and a consistent negative sign.

**Table 1:**
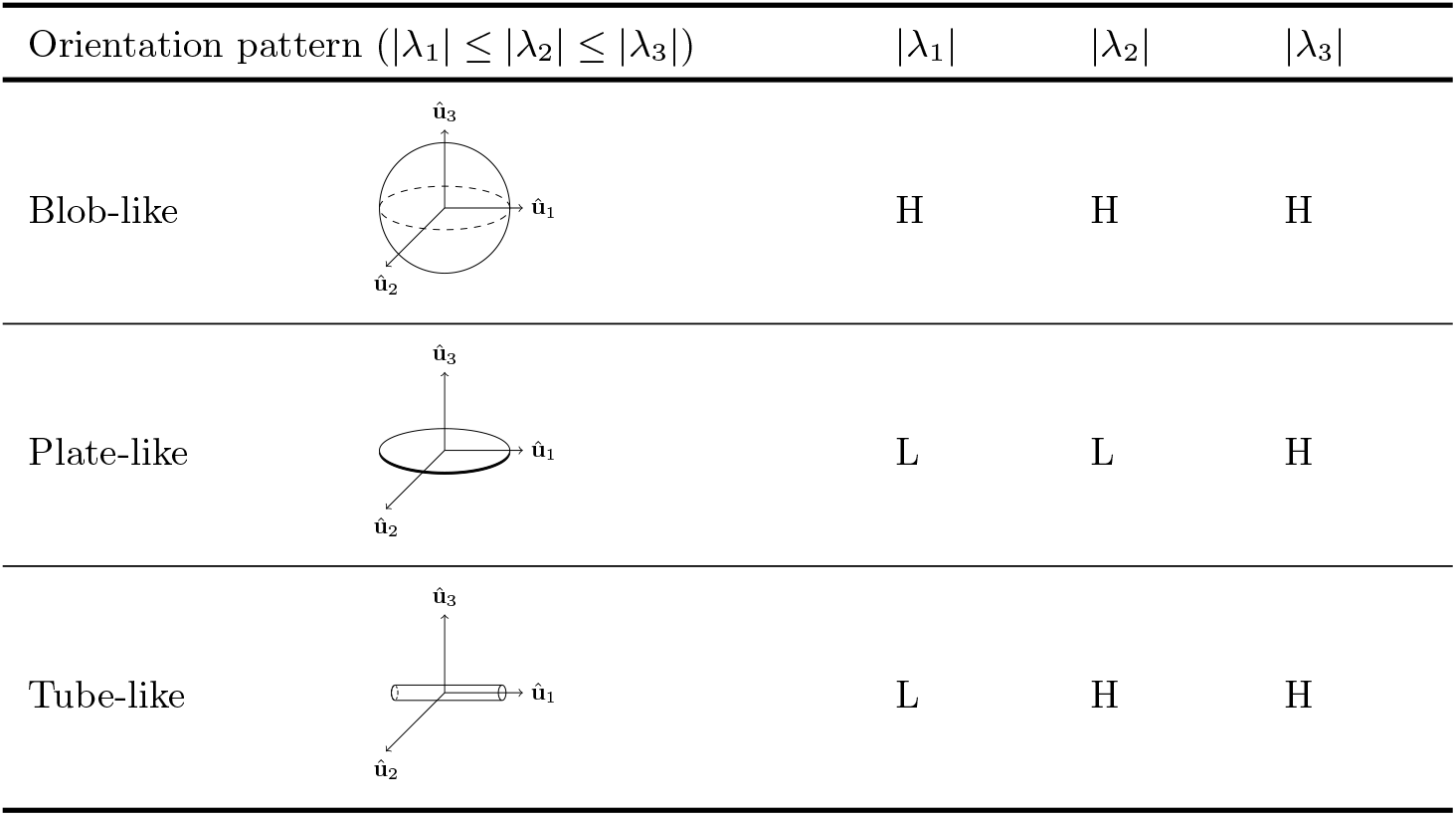
Relations between the eigenvalues of the Hessian matrix for different geometrical patterns (L/H: low/high absolute value; adapted from (Oruganti et al. (2013)). 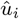 denote the normalized eigenvectors related to the i^th^ eigenvalue. Tubular structures are characterized by large-magnitude *λ*_2_ and *λ*_3_ eigenvalues and by a small dominant *λ*_1_; the sign of *λ*_2_ and *λ*_3_ is determined by the contrast polarity (negative for a positive contrast, i.e. light objects against a dark background).

| Orientation pattern ($\vert \lambda_1\vert \leq \vert \lambda_2\vert \leq \vert \lambda_3\vert $) | | $\vert \lambda_1\vert $ | $\vert \lambda_2\vert $ | $\vert \lambda_3\vert $ |
| --- | --- | --- | --- | --- |
| Blob-like |  | H | H | H |
| Plate-like |  | L | L | H |
| Tube-like |  | L | H | H |

On this basis, the first geometrical feature proposed by Frangi et al. accounts for the similarity to blob-like structures, which attains its maximum for locally isotropic objects (Table 1, first row):

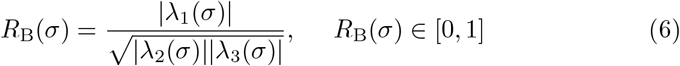

This *blobness* metric alone, however, would not discriminate between plate-shaped and tubular objects; accordingly, Frangi et al. introduced an additional geometrical grey-level-invariant feature, which evaluates the aspect ratio of the image ellipsoid in the plane orthogonal to the direction of minimum contrast, i.e. the ratio between the two largest second derivatives:

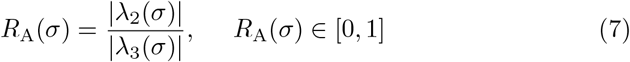

As shown in Table 1, this latter feature decays to 0 only within the neighborhood of plate-like structures, where |*λ*_2_(*σ*)| becomes significantly lower than |*λ*_3_(*σ*)|. Furthermore, Frangi et al. adopted a feature of image *structureness*, defined as the Frobenius norm of the Hessian matrix, which enables the distinction of low-contrast background regions, generally associated with small spatial derivatives and, thus, small eigenvalues:

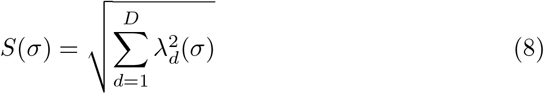

The above features are finally combined in a probability-like *vesselness* function which, in the case of a positive contrast polarity, is defined as:

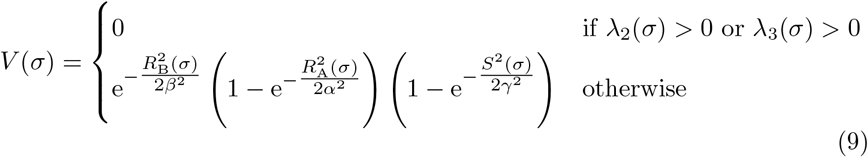

where the product of the basic exponential terms implies the suppression of all image regions which do not concurrently meet all the above criteria. *β, α* and *γ* are internal parameters of the Frangi filter which respectively tune its sensitivity to variations in the *R*_B_, *R*_A_ and *S* features. Namely, an increase in *β* would lead to a relatively higher sensitivity to blob-like structures, decreasing the related penalty introduced in the evaluation of the *vesselness* probability. On the other hand, lower *α* values would amplify the response of the filter to the presence of elongated structures. Usually, the *α* and *β* sensitivity parameters are heuristically fixed for the specific application or image modality of interest. Conversely, since the structureness *S* is directly influenced by the dynamic range of the raw input image, the related *γ* sensitivity may be automatically adapted: according to Frangi et al., setting *γ* to half of the maximum Hessian norm obtained at each scale (i.e., half of the maximum image structureness, *S*(*σ*)) ensures robust results.

The Frangi filter was devised as a multiscale method: accordingly, the above vesselness likelihood function is estimated for different levels of interest of the image scale-space, after a convolution with Gaussian kernels of different variance, *σ*^2^. As done in the present implementation of the filter, the multiscale analysis of the vesselness probability function may be parallelized over multiple cores, thus making the computational time independent from the number of spatial scales of interest. Finally, the maximum intensity projection (MIP) of the filter response along the image scale dimension is considered:

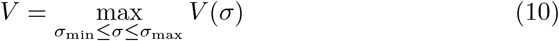

where, as discussed in the following, *V* (*σ*) locally reaches its maximum at a scale directly related to the cross-sectional diameter of the tubular structures present in the image. The main steps of the Frangi filter’s algorithm are summarized in Fig. 1.

**Figure 1:**
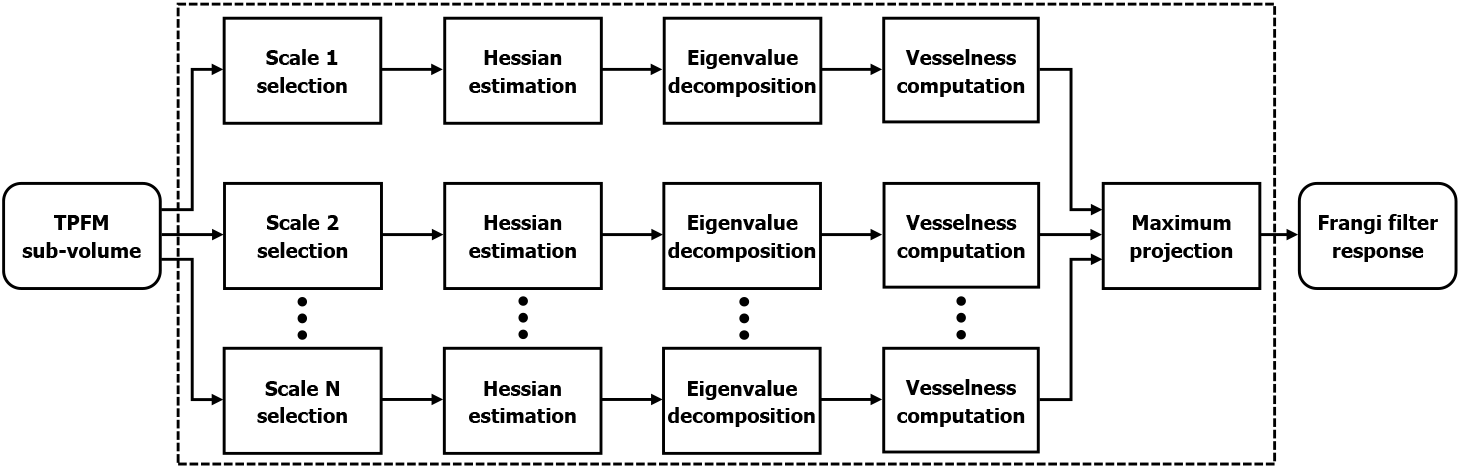
Block scheme representing the main computational steps of the Frangi filter. In the present application, the estimation of the *vesselness* probability function over different spatial scales was parallelized for efficiency’s sake.

### 2.4. Fiber orientation analysis pipeline

The image processing pipeline for the 3D analysis of myelinated fiber orientations was developed in Python and is made available as an open-source software at https://github.com/lens-biophotonics/Foa3D. The following paragraphs describe its main stages.

#### 2.4.1. TPFM image preprocessing

The separate TPFM image tiles composing the mesoscale reconstructions of the acquired brain slices are preliminarily aligned using ZetaStitcher, a custom software tool for high-resolution large volumetric stitching (Mazzamuto et al. (2018); Mazzamuto (2021)). The ZetaStitcher tool is also employed to programmatically access and iteratively process basic sub-volumes fitting the system’s memory, thus enabling the analysis of submicron-resolution TPFM reconstructions of arbitrary extension.

First, however, the uneven illumination of the separate TPFM stacks is adjusted via the retrospective CIDRE correction method (Smith et al. (2015)), to suppress the stitching artifacts which would arise when fusing the aligned TPFM tiles. The spatial intensity gain models related to the *λ*=618 nm and *λ*=482 nm wavelengths, shown in Fig. 2, were estimated using a reference dataset comprising 23118 and 66297 TPFM images, respectively.

**Figure 2:**
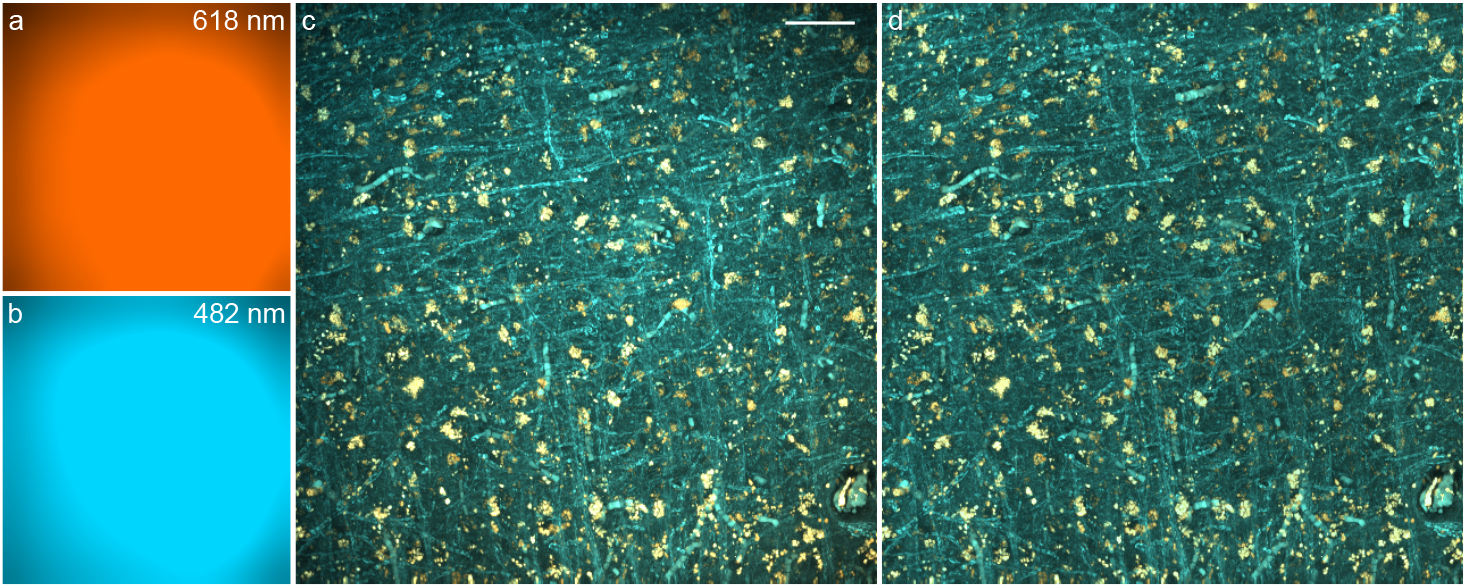
Example of the CIDRE-based tile illumination correction (MIPs): a) spatial intensity gain model at *λ*=618 nm (neurons); b) gain model at *λ*=482 nm (myelinated fibers); c) original TPFM tile; d) corrected TPFM tile (zero-light preserved mode (Smith et al. (2015)); scale bar: 50 μm.

Furthermore, if required, the coronal XY-plane of the sliced TPFM sub-volumes is preliminarily blurred using a 2D Gaussian smoothing filter, adopting a variance tailored with respect to the anisotropy of the PSF of the TPFM system; i.e., 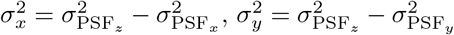, where 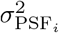 represents the variance of the PSF model along the i-axis. Indeed, confocal laser scanning and light-sheet microscopes provide 3D images characterized by a poorer resolution along the direction of the optical axis: if not properly corrected, the anisotropic resolution of the microscope would introduce a systematic bias in the computation of the Hessian matrices of the second directional derivatives and, thus, in the resulting 3D orientation estimates (Morawski et al. (2018)). Conversely, this transverse degradation of the optical resolution may be skipped if the TPFM tiles are preliminarily deconvolved. Following this optional smoothing stage, TPFM images are downsampled in the coronal plane, in order to obtain an isotropic 1 μm × 1 μm × 1 μm voxel size.

While the TPFM tile alignment and illumination correction must be performed separately via the respective tools, the image resolution isotropization is integrated within the developed pipeline.

#### 2.4.2. Multiscale Frangi filter

The Frangi filter was originally designed as a multiscale method for the analysis of angiography images (Frangi et al. (1998)), able to provide a concurrent enhancement of major arteries and veins, and peripheral blood vessels characterized by different cross-sectional sizes. However, for avoiding image artifacts, the analysis scales of the filter should be rigorously tuned with respect to the expected radius of the tubular structures of interest. As shown in (Oruganti et al. (2013)), the smoothing Gaussian kernel behaves similarly to a Dirac delta function if its standard deviation becomes too small compared to the diameter of the analyzed myelinated fibers; as a consequence, the scale-selection stage would yield the original fiber structure, and the subsequent analysis of the local Hessian matrices and estimation of the *vesselness* score would produce a spurious high-probability ridge at the fiber edge. Conversely, as the standard deviation gets relatively large, the Gaussian convolution would yield the kernel itself, with the fiber starting to act as a Dirac function. Fig. 3a shows the cross-sectional view of the Frangi’s vesselness probability of an ideal cylinder simulating a 3D myelinated axon (pixel density: 50 px*/*μm; length/diameter aspect-ratio: 12), related to different scale-to-radius ratios. The evaluation of the 1D vesselness profile (Fig. 3b) in terms of maximum relative intensity (Fig. 3c) and FWHM (Fig. 3d) shows half of the expected fiber radius to be the optimal spatial scale which best preserves the original intensity and cross-sectional size of the analyzed tubular structure.

**Figure 3:**
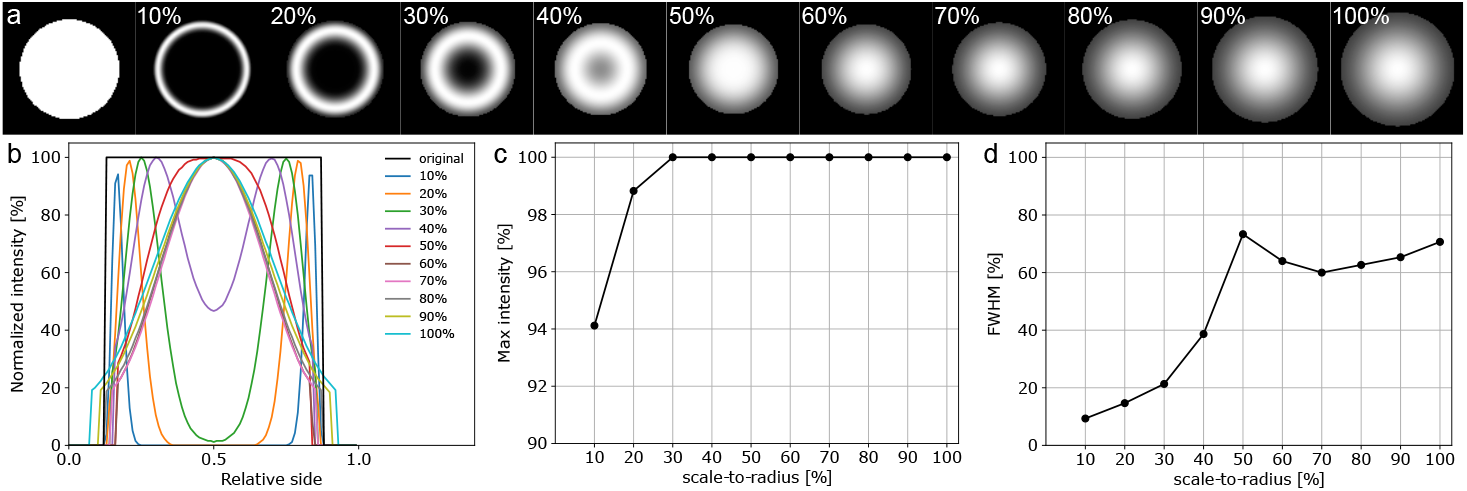
a) Cross-sectional view of the Frangi’s vesselness probability of a simulated 3D fiber (left), obtained by adopting increasing scale-to-radius ratios; b) 1D cross-sectional vesselness profiles, normalized with respect to the original intensity; c) normalized maximum intensity; d) normalized FWHM. The optimal scale corresponds to a scale-to-radius ratio of 50 %.

Regarding the three sensitivity parameters of the Frangi filter, *γ* was automatically tailored for each filtered sub-volume, and filtering scale, to the maximum norm of the estimated Hessian matrices (section 2.3). On the other hand *β* and *α*, which respectively tune the sensitivity of the filter to the geometrical features *R*_B_ and *R*_A_ (Eqs. 6, 7), were empirically adjusted by visually inspecting the myelinated fiber enhancement and the background suppression in the *λ*=482 nm channel, while exploiting the *λ*=618 nm channel as a comparative reference where neuronal bodies can be clearly distinguished due to the characteristic autofluorescence of lipofuscins, a fluorescent pigment that accumulates with ageing in the lysosomal compartment of postmitotic cells, including neurons (Moreno-García et al. (2018)). This evaluation led to a final sensitivity configuration given by *α* = 0.001 and *β* = 1 which, as made evident in Figs. 4a and 4b, is able to produce a marked selective enhancement of tubular fiber structures and a considerable rejection of the cell soma.

**Figure 4:**
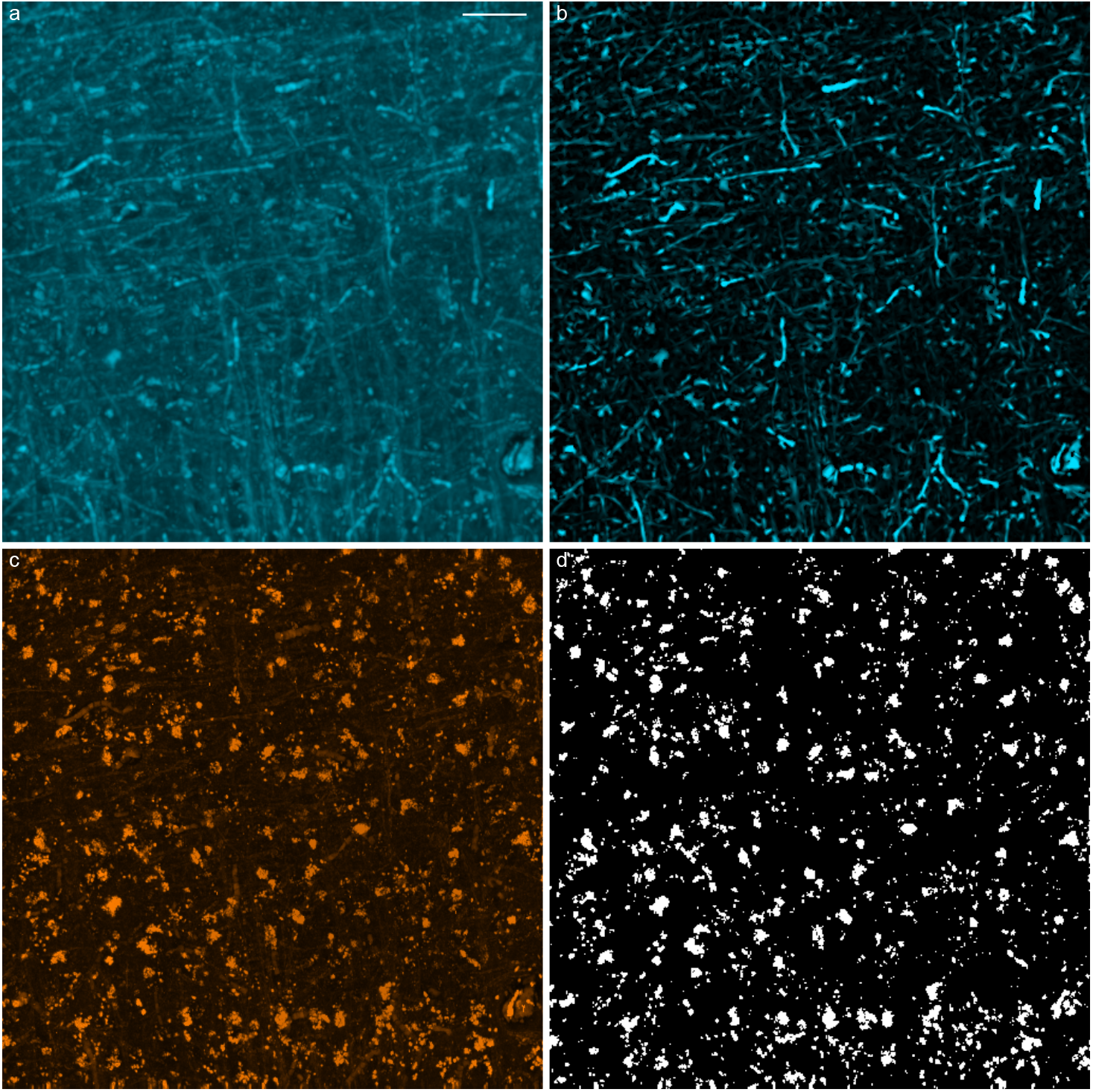
Frangi-based enhancement of myelinated fibers in the 3D TPFM tile shown in Fig. 2 (MIPs): a) myelinated fibers autofluorescence (*λ*=482 nm), enhanced via the MAGIC preparation protocol (Costantini et al. (2021)); b) Frangi’s fiber probability map (*α* = 0.001, *β* = 1, scales = (1, 1.25, 1.5) μm); c) neuronal bodies (*λ*=618 nm); d) neuron rejection mask (Yen’s thresholding method); scale bar: 50 μm.

**Figure 5:**
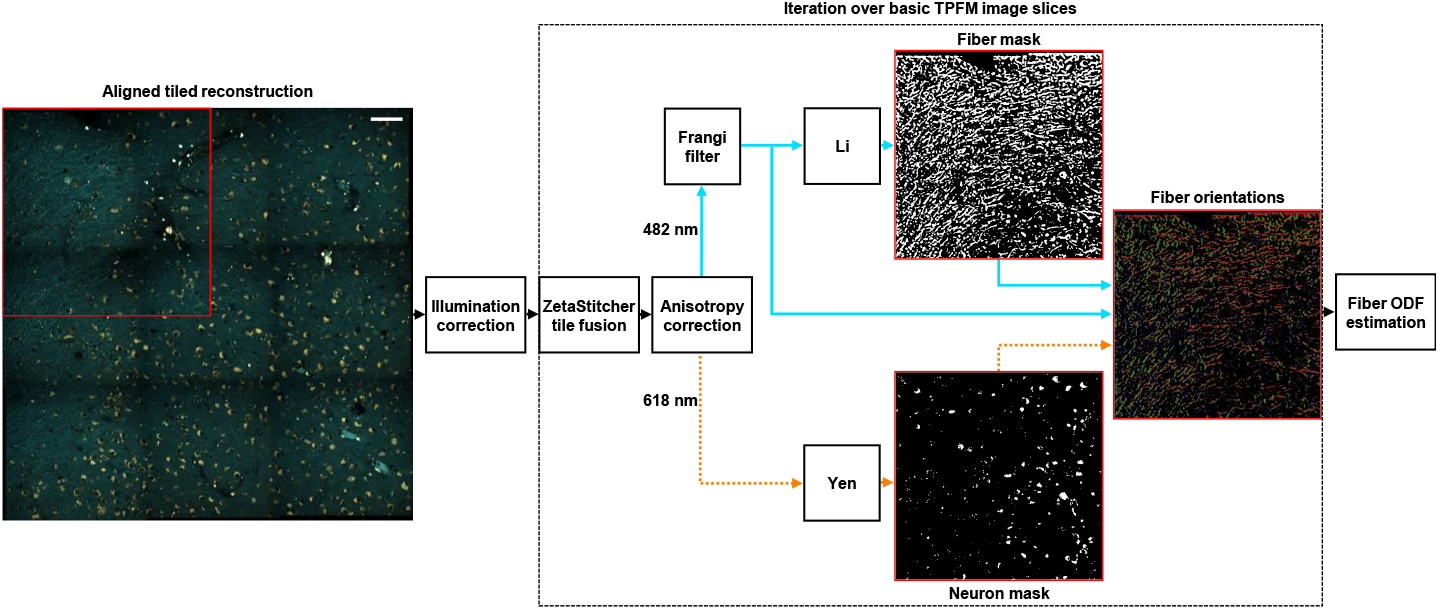
Block diagram schematizing the image processing pipeline for the 3D analysis of myelinated fiber orientations; the pipeline is iteratively applied to basic sub-volumes of the whole tiled reconstruction; scale bar: 100 μm.

In this work, the Frangi-enhanced images were binarized using Li’s minimum cross entropy thresholding method (Li and Tam (1998)), and 3D fiber orientations were finally identified as the eigenvectors associated with the dominant eigenvalues of the Hessian matrices belonging to voxels classified as myelinated fiber structures. In order to improve the specificity of the resulting fiber orientation maps, provided by the inherent attenuation of non-tubular objects achieved by the Frangi filter, the developed pipeline optionally performs a postprocessing step which suppresses the neuronal bodies by masking the *λ*=618 nm channel via the Yen’s binarization algorithm (Jui-Cheng Yen et al. (1995)). An example of this lipofuscin autofluorescence-based neuron segmentation is shown in Figs. 4c and 4d.

#### 2.4.3. Analytical fiber Orientation Distribution Functions

High-resolution fiber orientation data obtained at the native pixel size of the TPFM system can be integrated into Orientation Distribution Functions (ODFs) (Bunge (1982)), providing a comprehensive statistical description of 3D fiber tract orientations within larger spatial compartments or super-voxels. Due to their widespread use in the neuroimaging community, the estimation of TPFM-based ODFs is highly suitable for a multimodal quantitative comparison with spatial fiber architectures mapped by other advanced optical modalities, as 3D-Polarized Light Imaging (Axer et al. (2011)). Furthermore, the spatial downscaling produced by the ODF estimation allows to bridge the gulf between single fiber orientations obtained via optical microscopy and the meso- and macro-scale connectomics that is generally targeted by diffusion magnetic resonance imaging (Axer et al. (2016)). In the developed image processing pipeline, the ODFs of myelinated fibers were estimated from the 3D orientation vector fields returned by the Frangi filtering stage by means of the fast analytical approach proposed by Alimi (Alimi et al. (2018, 2019)). The present method is computationally efficient and is characterized by improved angular precision and resolution with respect to deriving the ODFs by modeling local directional histograms of discretized fiber orientations (Alimi et al. (2020)). In detail, fiber orientation vectors are parametrized in terms of their coordinates *ϕ* and *θ* = 90^°^ − *α* on the unit sphere *S*^2^, which respectively denote the azimuth angle representing the orientation of the fiber axis within the XY-plane, and the polar angle related to the *α* angle between the fiber and the XY-plane. Their square-integrable statistical distribution *f* (*θ, ϕ*) ∈ *L*^2^(*S*^2^) can then be expanded as a linear combination of spherical harmonics:

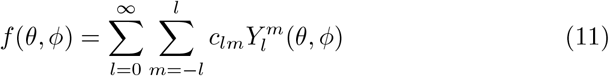

where 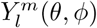 are the spherical harmonics of order *l* and degree *m*, defined by:

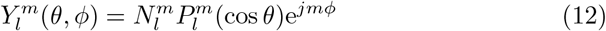

In the above expression, 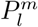 represents the associated Legendre polynomial whereas 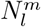 is a normalization factor given by:

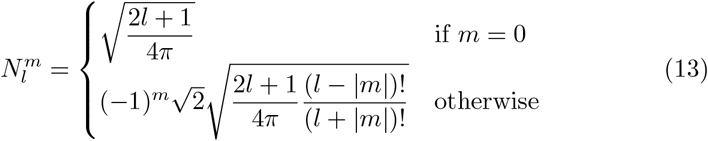

The coefficients *c*_*lm*_ uniquely describing the fiber orientation distribution *f* (*θ, ϕ*) can be recovered from its spherical Fourier transform, defined as:

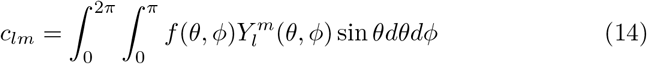

In the adopted approach, fiber orientation vectors are regarded as two-dimensional Dirac delta functions on *S*^2^; thus, the distribution *f*_SV_(*θ, ϕ*) of *K* orientations included in a given super-voxel can be modeled as follows:

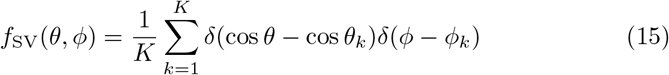

As shown by Alimi et al., substituting Eq. 12 and Eq. 15 into Eq. 14, and applying the sifting property of Dirac functions, finally yields the continuous analytical expression of the spherical harmonics coefficients, reported below:

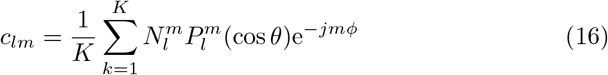

In this work, the series expansion of the analytical ODF was limited to the 6^th^ order (*l*_max_=6), as previously performed by Axer et al. for the estimation of 3D-PLI-based fiber ODFs (Axer et al. (2016)).

#### 2.4.4. Pipeline configuration and computational runtimes

The runtime performance of the designed image processing pipeline was evaluated on a high-performance computer cluster comprising a pair of twin computing nodes, each equipped with 192 GB of RAM and two Intel^®^ Xeon^®^ Silver 4214R 12-core processors operating at 2.4 GHz.

A first evaluation concerned the variation in the computational time and the memory usage of the Frangi-based fiber orientation analysis stage with respect to the number of spatial scales employed by the Frangi filter. In order to ensure an unbiased comparison, a fixed spatial scale of 1.25 μm was adopted in this regard i.e., a different number of scales of identical value was fed to the pipeline. Furthermore, the execution time of the entire processing workflow, including the generation of the Frangi-based fiber ODFs, was assessed adopting different combinations of the size of the basic image chunks analyzed iteratively, and the size of the orientation map super-voxels defined for the estimation of the fiber ODFs. In detail, chunk sizes of 50, 100, 150, 200 and 250 MB, and ODF super-voxel sides of 16, 32, 48, 64 and 80 μm were tested in this work. The present tests were conducted on a reference 8859.9 μm × 5204.8 μm × 98.0 μm TPFM reconstruction of a human brain slice (252 3D tiles, 16.4 GB).

#### 2.4.5. Validation of fiber orientation estimates

An *ad hoc* validation module was developed for verifying the reliability of the 3D orientation estimates returned by the image processing pipeline designed in this work. 10 TPFM image stacks acquired including grey matter (GM) tissue and 10 stacks including white matter (WM) were independently considered. The open Zenodo dataset including these image stacks can be accessed at: https://doi.org/10.5281/zenodo.7181945. Based on the characterization of the scale-dependent response of the Frangi filter to tubular structures of different cross-sectional size (section 2.4.2), three spatial scales, namely 1, 1.25 and 1.5 μm, were adopted in the present validation; these respectively correspond to an optimal fiber diameter of 4, 5 and 6 μm. In detail, an automatic image patch generator was conceived so as to randomly sample and analyze basic 75 μm×75 μm×15 μm (x, y, z) slices of the original TPFM stacks. The ranges of the sliced sub-volumes were properly extended in order to cope with the boundary artifacts caused by the convolution with the smoothing Gaussian kernel, implementing the scale selection step of the Frangi filter. Furthermore, each sampled image patch was randomly flipped along its three axes with the aim to decrease the dependency of the obtained results on the orientation statistics of the assessed image stacks. In detail, similarly to the approach followed in (Giardini et al. (2021)), the sliced patches were separately rotated about the z- and x-axis over a −45^°^ to 45^°^ range, with a sampling step of 5^°^, in order to produce a controlled reorientation of myelinated fiber structures, respectively within and outside of the image plane of the TPFM system. A bicubic spline interpolation with prefiltering was applied when rotating the randomly sampled patches. The transformed images, and their original, unrotated version, were separately processed by the developed pipeline in order to derive patch-wise distributions of the in-plane azimuth angle *ϕ*_*xy*_ and the out-of-plane elevation angle *θ*_*zy*_. At this stage, the extended edge introduced to account for the smoothing-related boundary artifacts was properly masked prior to the evaluation of the angular distributions. These were corrected according to the applied 1D rotations, i.e. rotating the angular reference system, and then compared to the distributions generated from the original image. Specifically, the discrepancy between the distributions was characterized by means of the inter-median distances 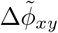 and 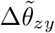 and, secondly, by their Bhattacharyya coefficient (Bhattacharyya (1946)):

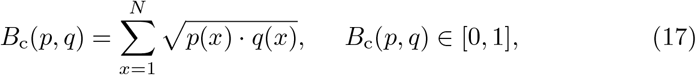

which estimates the amount of overlap between two statistical samples, computed here adopting N=180 bins of 1^°^ in width. A threshold of 1 % was imposed on the relative number of pixels classified as belonging to myelinated fibers: TPFM patches not meeting this criterion for each tested rotation were systematically excluded from further consideration. This ensured that the above metrics were obtained only from adequately populated data samples, with an equal number of patches related to each tested rotation value.

## 3. Results

In order to validate the reliability of the estimated 3D fiber orientation maps, random image slices automatically extracted from a reference dataset of 10 GM and 10 WM stacks were methodically rotated within and out of the image plane of the TPFM system, and fed to the Frangi-based orientation analysis pipeline, quantitatively evaluating the degree of agreement between the resulting distributions of the fiber azimuth (*ϕ*_*xy*_) and elevation (*θ*_*zy*_) angles and the ones obtained from the original unrotated slices, in terms of their median value and their Bhattacharyya coefficient. The results of the validation procedure are summarized in Fig. 6, which separately shows the accuracy performance related to the assessment of the *ϕ*_*xy*_ and *θ*_*zy*_ angles in GM and WM image samples. A detailed example of the adopted validation strategy is provided in the Supplementary Materials (Fig. **S1**).

**Figure 6:**
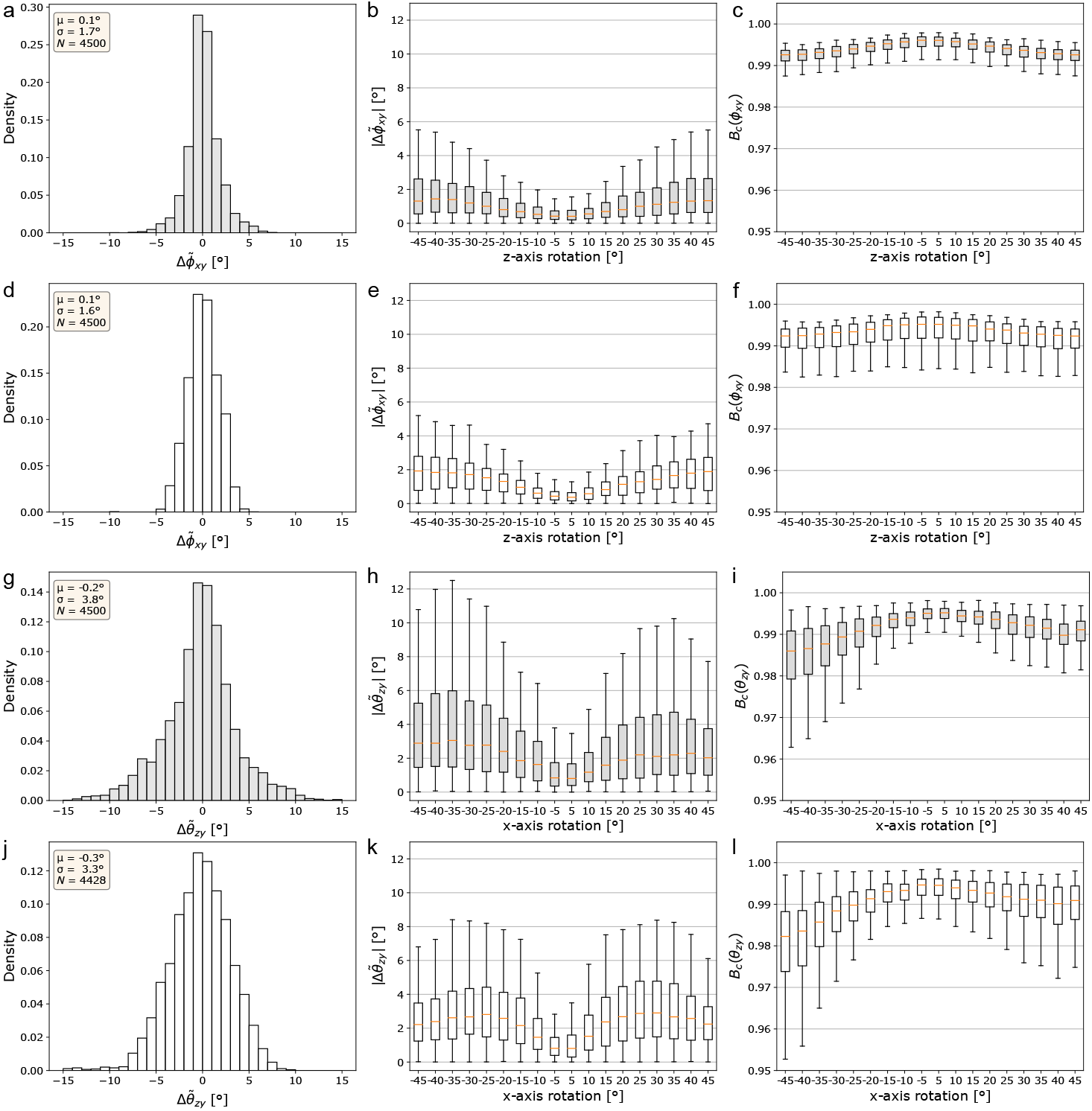
Validation of the 3D Frangi-based fiber orientation maps accuracy (*N* = *N*_TPFM_ *× N*_ROT_, where *N*_TPFM_ denotes the total number of sample image patches, and *N*_ROT_ = 18 is the number of test rotations). Left column: overall distributions of the inter-median errors; center column: absolute inter-median distances against the test rotation applied; right column: Bhattacharyya coefficient against test rotations. a, b, c) azimuth angle *ϕ*_*xy*_, grey matter images; d, e, f) azimuth angle *ϕ*_*xy*_, white matter; g, h, i) elevation angle *θ*_*zy*_, grey matter; g, h, i) elevation angle *θ*_*zy*_, white matter.

In specific, the overall distributions grouping the inter-median errors pertaining to each tested rotation range (Figs. 6a, 6d, 6g, 6j) exhibit a near-zero mean value, which implies the absence of a systematic bias in the developed fiber orientation analysis. However, their standard deviation is notably lower in the case of the *ϕ*_*xy*_ orientations evaluated within the imaging plane of the TPFM system, where the optical resolution achieved is higher. This is made evident by the trend of the absolute inter-median error 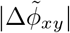 along the whole range of tested rotations (Figs. 6b, 6e) whose median value, despite exhibiting a sensible increase for larger rotations, never exceeds 2^°^ in the case of the azimuth angular coordinate *ϕ*_*xy*_, in both the test sets of GM and WM image patches. Conversely, Figs. 6h and 6k show a tendency towards relatively larger intermedian distances 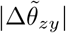 for the elevation angle between the myelinated fiber structures detected by the pipeline and the imaging plane of the TPFM system. Despite being higher, median error values nevertheless stay below 3^°^ in both the evaluated GM and WM samples. Moreover, as highlighted by the boxplots of the Bhattacharyya coefficient, representing the amount of overlap with the original reference angular distributions of *θ*_*zy*_ (Figs. 6i, 6l), the image processing pipeline appears to exhibit an asymmetrical performance with respect to the direction of the applied rotation. This may be ascribed to the preferential orientation statistics of the selected WM image tiles, despite the random axis inversion preliminarily applied to the image patches considered in the present validation. On the other hand, consistently with the trend of the inter-median distance 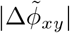, the estimated Bhattacharyya coefficients are remarkably uniform for the different planar rotations applied about the z-axis, with their median value never dropping below 0.99 in both the GM and WM image samples (Figs. 6c, 6f). Table **S1** and **S2** in the Supplementary Materials report the complete median values and interquartile ranges of the distributions shown in Fig. 6.

Fig. 7 summarizes the runtime performance of the Frangi-based fiber orientation analysis pipeline, applied to the TPFM reconstruction shown in Fig. 8. In detail, using Frangi filters with an increasing number of spatial scales determines an expected rise in the memory usage of the fiber orientation analysis stage (Fig. 7a). However, by parallelizing the estimation of the vesselness probability function (Eq. 9, Fig. 1), the interrogation of multiple scales does not lead to a proportional increase in the computational time of the filter. This accounts for a large percentage of the total execution time of the pipeline (Fig. 7b), with respect to the subsequent estimation of the fiber ODFs (Fig. 7c). As shown in (Alimi et al. (2020)), the adoption of larger ODF super-voxel sizes, and the consequent decrease in their overall number, is not accompanied by a monotonic reduction in the time required to generate the fiber ODF maps of the whole TPFM image volume: this is related to the continuous setting of Alimi’s analytical approach, which computes the spherical harmonics coefficients (16) for each native voxel of the ODF compartments, independently from their size. Similarly, processing larger chunks of the TPFM reconstruction does not appear to be generally beneficial for the computational efficiency of the developed processing pipeline, with a tendency towards longer execution times when the basic chunk size is increased, in spite of their lower number (Fig. 7b). Fig. 8 finally highlights the capability of the image processing pipeline to reproduce the orientation of myelinated fiber structures at multiple spatial scales in unstained brain tissue slices, imaged with TPFM. Adopting differently sized ODF super-voxels, thus producing a different level of downscaling of the high-resolution vector fields returned by the Frangi-based orientation analysis, allows to shift the focus from microscopic single fiber tracts to mesoscopic long-range fiber bundles in white matter areas.

**Figure 7:**
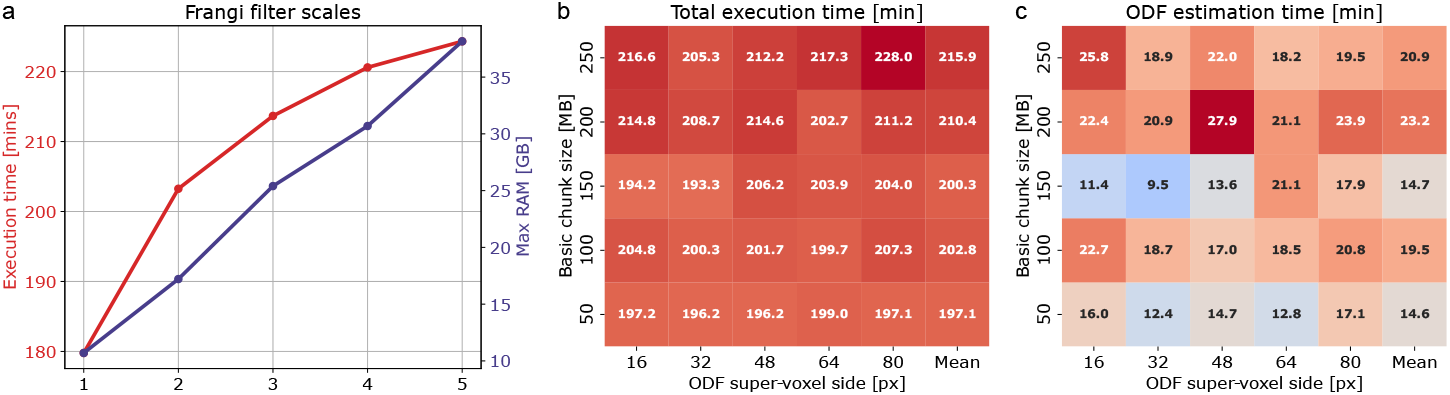
Runtime performance of the image processing pipeline: a) execution times and memory usage of the Frangi-based fiber orientation analysis stage against the number of spatial scales of the Frangi filter (*σ* = 1.25 μm, image chunk size = 50 MB); b) total execution time against different combinations of the spatial resolution of the fiber ODFs and the size of the basic image chunks analyzed iteratively (*σ* = 1.25 μm); fiber ODFs estimation time, against different configurations of the designed pipeline.

**Figure 8:**
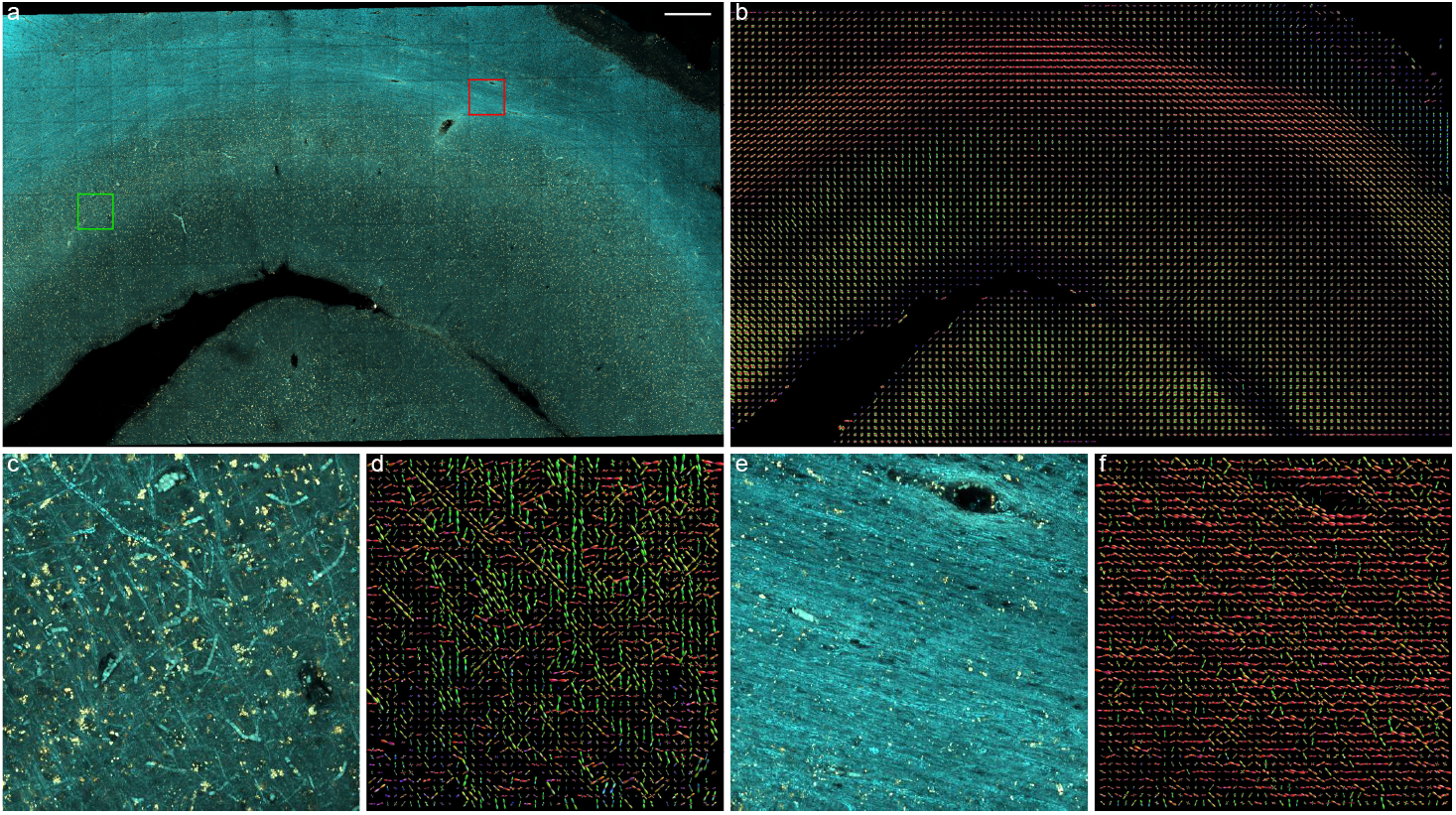
Frangi-based orientation analysis of myelinated fibers in a mesoscale TPFM reconstruction of a portion of a human brain 60 μm-thick slice: a) original tiled reconstruction aligned and fused using ZetaStitcher (scale bar: 0.5 mm) (Mazzamuto et al. (2018); Mazzamuto (2021)), following the uneven illumination correction achieved via the CIDRE method (Smith et al. (2015)); b) myelinated fiber ODF map of the TPFM reconstruction returned by the image processing pipeline (*α* = 0.001, *β* = 1, scales = (1.25, 2.5) μm; ODF super-voxel: 80 μm *×* 80 μm *×* 80 μm); c) MIP of a 10 μm-deep section including grey matter (green square, scale bar: 50 μm); d) corresponding grey matter ODFs (super-voxel: 10 μm *×* 10 μm *×* 10 μm); MIP of a 10 μm-deep section including white matter (red square); f) corresponding white matter ODFs (super-voxel: 10 μm *×* 10 μm *×* 10 μm). Fiber ODF maps were displayed using the open-source software package for medical image processing and visualisation MRtrix3 (Tournier et al. (2019)).

## 4. Discussion

In this study, we employed two-photon excitation microscopy with the glycerol-based MAGIC tissue preparation protocol to acquire 3D micron-resolution mesoscopic reconstructions of the human brain myeloarchitectonics, without the requirement of exogenous staining of myelinated fiber axons. We then applied the image processing tool presented in this work to derive fiber-specific orientation maps, exploiting a multiscale 3D Frangi filter for obtaining a targeted enhancement and segmentation of elongated image structures. The optimal spatial scale of the filter with respect to the cross-sectional size of myelinated fibers was thoroughly assessed, whereas the internal sensitivity parameters related to the geometrical features exploited by the enhancement mechanism were empirically tailored through a qualitative evaluation of the improvement of the fiber autofluorescence contrast at 482 nm, and the suppression of neuronal bodies with respect to the soma-specific fluorescence emission at 618 nm. A data-driven fine-tuning of these parameters on a manually generated ground truth of classified TPFM images of myelinated fiber structures may be considered in future work. Nonetheless, the results of the validation procedure outlined in section 2.4.5 demonstrate that the current configuration of the fiber orientation analysis pipeline is able to provide an accurate evaluation of white matter fiber tract and grey matter single fiber 3D orientations, even without the application of cutting-edge deep learning techniques, and despite the widening of the point-spread-function along the optical axis of the employed microscopy system. In general, the poorer axial resolution of fluorescence microscopes, LSFM systems particularly, poses a critical restraint on the feasibility of a 3D quantification of myelinated fiber orientations, and has forced other authors to limit their studies to 2D analyses orthogonal to the optical axis (Budde et al. (2011); Morawski et al. (2018)). In the present work, this translated in a relatively higher reliability of the *ϕ*_*xy*_ fiber azimuth angles, both in grey and white matter regions. The achieved performances may, however, increase as a result of a preliminary deconvolution of the TPFM image volumes, not applied here because of the severe spatial undersampling of the PSF. Nevertheless, as pointed out by Morawski (Morawski et al. (2018)), this inherent limitation can be overcome by employing multi-view imaging systems, such as dual inverted selective plane illumination microscopes (diSPIM) (Kumar et al. (2014)), and making use of dedicated multi-view deconvolution algorithms.

The unsupervised nature of the developed fiber orientation analysis tool brings the key advantage of relaxing the need for a high-quality ground truth of volumetric segmentations of fiber architectures, manually annotated by expert operators. Nevertheless, cascading the 3D Frangi filter with state-of-the-art deep neural networks, such as the 3D U-Net classifier recently proposed in (Ç içek et al. (2016)) may lead to a remarkable improvement of the automatic identification of myelinated fibers with respect to the binarization mechanism of Li’s thresholding algorithm, which minimizes the cross entropy between the classified foreground and the foreground mean, and the background and the background mean. Training 3D deep learning models generally requires a comprehensive manual annotation of volumetric microscopy images, possibly entailing an overwhelming human effort that was not feasible in the present work. However, in this regard, semi-supervised frameworks as the one introduced in (Lee (2013)), which make use of sparse partial annotations complemented by pseudo-labels inferred on unlabelled regions, may speed up the training process.

In the present study, the accuracy of the Frangi-based fiber orientation analysis pipeline was evaluated at the micron scale, analyzing image slices randomly sampled from a set of TPFM stacks of grey and white matter, and properly rotated within and out of the image plane of the microscopy setup. In view of this, future work may consider a multimodal evaluation of myelinated fiber orientations, comparing the 3D vector maps returned by the developed pipeline with the ones directly generated by means of polarimetry-based techniques, such as RP-CARS (Rotating-Polarization Coherent Anti-Stokes Raman Scattering) (Cheng et al. (2001); de Vito et al. (2012)) and the already mentioned 3D-PLI and 3D-PSOCT. Indeed, the unsurpassed spatial resolution of fluorescence microscopy may help corroborating 3D-PLI and 3D-PSOCT fiber orientation estimates or, on the other hand, characterize potential limitations of these polarimetry-based modalities.

Most importantly, the combination of the label-free MAGIC preparation protocol, TPFM, and 3D Frangi-based analysis of myelinated fiber orientations may represent a powerful approach for a quantitative histological validation of 3D fiber ODF maps obtained by dMRI, aiding the assessment of the brain region-specific reliability of state-of-the-art dMRI-based tractography.

## Author Contributions

**Michele Sorelli**: Writing – original draft, Software, Conceptualization, Investigation, Formal analysis, Visualization, Writing - review & editing. **Irene Costantini**: Conceptualization, Investigation, Resources, Supervision, Project administration, Writing - review & editing. **Leonardo Bocchi**: Supervision, Writing - review & editing. **Markus Axer**: Resources, Writing - review & editing. **Francesco Saverio Pavone**: Resources, Funding acquisition, Writing - review & editing. **Giacomo Mazzamuto**: Resources, Supervision, Software, Writing - review & editing.

