## Supplementary Material for "Fiber enhancement and 3D orientation analysis in label-free two-photon fluorescence microscopy"

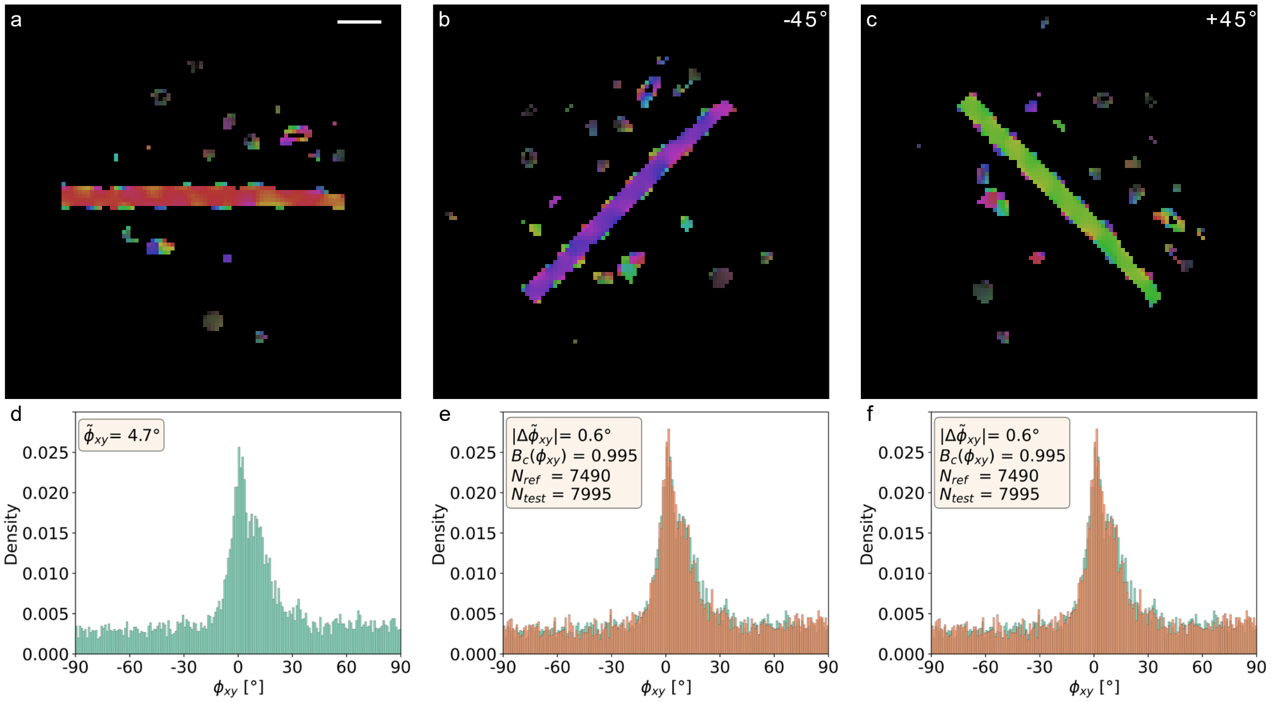


**Figure S1** Example of the test rotations applied for validating the accuracy of the Frangi-based fiber orientation analysis pipeline (α = 0.001, β = 1, scales = [1, 1.25, 1.5] µm); the sliced 75 µm x 75 µm x 15 µm grey matter patch includes a myelinated fiber having an approximate diameter of ~5 µm: a) reference fiber orientation vectors, color-coded with respect to the in-plane azimuth angle $\varphi$ (scale bar: 10 µm); b, c) orientation vectors obtained following 1D rotations of -45° and +45° about the z-axis; d) original distribution of the fiber azimuth angles estimated at a 1 µm x 1 µm x 1 µm voxel size; e, f) angular distributions generated from the rotated image patches (orange), and expressed in the adjusted spatial reference system.

**Table S1** Median ($q_{0.5}$) and quartiles ($q_{0.25,\,0.75}$) of the inter-median distance between the original reference distributions of the azimuth $\varphi_{xy}$ and elevation $\theta_{zy}$ of myelinated fibers, and the (corrected) angular distributions generated following different 1D rotations around the z and x axes. The results related to grey (GM) and white matter (WM) image patches, randomly sampled from a test set of 3D TPFM tiles, are shown separately.

| Tissue | Rotation [°] | | -45 | -40 | -35 | -30 | -25 | -20 | -15 | -10 | -5 | +5 | +10 | +15 | +20 | +25 | +30 | +35 | +40 | +45 |
|---|---|---|---|---|---|---|---|---|---|---|---|---|---|---|---|---|---|---|---|---|
| $\Delta\tilde{\varphi}_{xy}$ | GM N=250 | $q_{0.75}$ | 2.6 | 2.5 | 2.4 | 2.2 | 1.8 | 1.5 | 1.2 | 1.0 | 0.7 | 0.8 | 0.9 | 1.2 | 1.6 | 1.8 | 2.1 | 2.4 | 2.6 | 2.6 |
| | | $q_{0.5}$ | 1.3 | 1.4 | 1.4 | 1.2 | 1.0 | 0.8 | 0.7 | 0.5 | 0.4 | 0.4 | 0.6 | 0.7 | 0.8 | 1.0 | 1.1 | 1.2 | 1.3 | 1.3 |
| | | $q_{0.25}$ | 0.6 | 0.7 | 0.6 | 0.6 | 0.6 | 0.4 | 0.3 | 0.3 | 0.2 | 0.2 | 0.3 | 0.4 | 0.4 | 0.4 | 0.5 | 0.6 | 0.6 | 0.6 |
| | WM N=250 | $q_{0.75}$ | 2.8 | 2.7 | 2.7 | 2.4 | 2.1 | 1.7 | 1.4 | 0.9 | 0.7 | 0.6 | 0.9 | 1.3 | 1.6 | 1.9 | 2.2 | 2.5 | 2.6 | 2.7 |
| | | $q_{0.5}$ | 1.9 | 1.8 | 1.8 | 1.7 | 1.5 | 1.3 | 1.0 | 0.6 | 0.4 | 0.4 | 0.6 | 0.8 | 1.1 | 1.3 | 1.4 | 1.7 | 1.8 | 1.9 |
| | | $q_{0.25}$ | 0.8 | 0.9 | 0.9 | 0.9 | 0.8 | 0.7 | 0.6 | 0.3 | 0.2 | 0.2 | 0.2 | 0.5 | 0.5 | 0.6 | 0.8 | 0.9 | 0.9 | 0.8 |
| $\Delta\tilde{\theta}_{zy}$ | GM N=250 | $q_{0.75}$ | 5.3 | 5.8 | 6.0 | 5.4 | 5.1 | 4.4 | 3.6 | 3.0 | 1.7 | 1.7 | 2.3 | 3.2 | 4.0 | 4.4 | 4.6 | 4.7 | 4.3 | 3.7 |
| | | $q_{0.5}$ | 2.9 | 2.9 | 3.0 | 2.8 | 2.8 | 2.4 | 1.9 | 1.6 | 0.8 | 0.8 | 1.2 | 1.6 | 1.9 | 2.2 | 2.1 | 2.2 | 2.3 | 2.0 |
| | | $q_{0.25}$ | 1.5 | 1.5 | 1.5 | 1.3 | 1.2 | 1.2 | 0.9 | 0.7 | 0.4 | 0.4 | 0.6 | 0.7 | 0.8 | 0.8 | 1.0 | 1.0 | 1.1 | 1.0 |
| | WM N=246 | $q_{0.75}$ | 3,5 | 3.7 | 4.2 | 4.3 | 4.4 | 4.1 | 3.8 | 2.6 | 1.5 | 1.6 | 2.8 | 3.8 | 4.5 | 4.8 | 4.8 | 4.6 | 3.9 | 3.3 |
| | | $q_{0.5}$ | 2.2 | 2.4 | 2.6 | 2.7 | 2.8 | 2.6 | 2.2 | 1.5 | 0.8 | 0.8 | 1.5 | 2.4 | 2.7 | 2.9 | 2.9 | 2.7 | 2.6 | 2.2 |
| | | $q_{0.25}$ | 1.2 | 1.3 | 1.4 | 1.6 | 1.5 | 1.3 | 1.1 | 0.7 | 0.4 | 0.3 | 0.7 | 0.9 | 1.2 | 1.4 | 1.5 | 1.4 | 1.3 | 1.3 |

**Table S2** Median ($q_{0.5}$) and quartiles ($q_{0.25,\,0.75}$) of the Bhattacharyya coefficient estimating the degree of overlap between the reference angular distributions of the azimuth $\varphi_{xy}$ and elevation $\theta_{zy}$ of myelinated fibers, and the (corrected) angular distributions generated following different 1D rotations around the z and x axes. The results related to grey (GM) and white matter (WM) image patches, randomly sampled from a test set of 3D TPFM tiles, are shown separately.

| **Tissue** | **Rotation [°]** | | **-45** | **-40** | **-35** | **-30** | **-25** | **-20** | **-15** | **-10** | **-5** | **+5** | **+10** | **+15** | **+20** | **+25** | **+30** | **+35** | **+40** | **+45** |
|---|---|---|---|---|---|---|---|---|---|---|---|---|---|---|---|---|---|---|---|---|
| $B_c(\varphi_{xy})$ | GM N=250 | $q_{0.75}$ | 0.994 | 0.994 | 0.994 | 0.995 | 0.995 | 0.996 | 0.996 | 0.997 | 0.997 | 0.997 | 0.997 | 0.996 | 0.996 | 0.995 | 0.995 | 0.994 | 0.994 | 0.994 |
| | | $q_{0.5}$ | 0.993 | 0.993 | 0.993 | 0.994 | 0.994 | 0.995 | 0.995 | 0.996 | 0.996 | 0.996 | 0.996 | 0.995 | 0.995 | 0.994 | 0.994 | 0.993 | 0.993 | 0.993 |
| | | $q_{0.25}$ | 0.991 | 0.991 | 0.992 | 0.992 | 0.993 | 0.993 | 0.994 | 0.994 | 0.995 | 0.995 | 0.995 | 0.994 | 0.993 | 0.993 | 0.992 | 0.992 | 0.991 | 0.991 |
| | WM N=250 | $q_{0.75}$ | 0.994 | 0.994 | 0.994 | 0.995 | 0.995 | 0.996 | 0.996 | 0.997 | 0.997 | 0.997 | 0.997 | 0.996 | 0.996 | 0.995 | 0.995 | 0.995 | 0.994 | 0.994 |
| | | $q_{0.5}$ | 0.992 | 0.992 | 0.993 | 0.993 | 0.993 | 0.994 | 0.995 | 0.995 | 0.995 | 0.995 | 0.995 | 0.995 | 0.994 | 0.994 | 0.993 | 0.993 | 0.993 | 0.992 |
| | | $q_{0.25}$ | 0.990 | 0.989 | 0.990 | 0.990 | 0.990 | 0.991 | 0.991 | 0.992 | 0.992 | 0.992 | 0.992 | 0.991 | 0.991 | 0.991 | 0.990 | 0.990 | 0.989 | 0.989 |
| $B_c(\theta_{zy})$ | GM N=250 | $q_{0.75}$ | 0.991 | 0.991 | 0.992 | 0.993 | 0.994 | 0.994 | 0.995 | 0.995 | 0.996 | 0.996 | 0.996 | 0.996 | 0.995 | 0.995 | 0.994 | 0.994 | 0.992 | 0.993 |
| | | $q_{0.5}$ | 0.986 | 0.987 | 0.988 | 0.989 | 0.991 | 0.992 | 0.994 | 0.994 | 0.995 | 0.995 | 0.994 | 0.994 | 0.994 | 0.993 | 0.992 | 0.991 | 0.990 | 0.991 |
| | | $q_{0.25}$ | 0.979 | 0.980 | 0.982 | 0.985 | 0.987 | 0.989 | 0.992 | 0.992 | 0.994 | 0.994 | 0.993 | 0.992 | 0.991 | 0.990 | 0.989 | 0.989 | 0.988 | 0.988 |
| | WM N=246 | $q_{0.75}$ | 0.988 | 0.988 | 0.990 | 0.992 | 0.993 | 0.994 | 0.995 | 0.995 | 0.996 | 0.996 | 0.996 | 0.996 | 0.995 | 0.995 | 0.995 | 0.995 | 0.994 | 0.994 |
| | | $q_{0.5}$ | 0.982 | 0.984 | 0.986 | 0.988 | 0.990 | 0.991 | 0.993 | 0.993 | 0.995 | 0.995 | 0.994 | 0.993 | 0.993 | 0.992 | 0.991 | 0.991 | 0.990 | 0.991 |
| | | $q_{0.25}$ | 0.974 | 0.975 | 0.980 | 0.983 | 0.986 | 0.988 | 0.991 | 0.991 | 0.992 | 0.992 | 0.991 | 0.991 | 0.989 | 0.988 | 0.987 | 0.987 | 0.985 | 0.986 |

**Video S1** Example video of a 3D fiber orientation colormap (x-component: red channel; y-component: green channel; z-component: blue channel) generated from a 14 μm-deep TPFM section (isotropized pixel size; original uneven illumination).
